# Data from an electronic health informatics pipeline to describe clearance dynamics of Hepatitis B surface antigen (HBsAg) and e-Antigen (HBeAg) in chronic HBV infection

**DOI:** 10.1101/494500

**Authors:** Louise O Downs, David Smith, Sheila F Lumley, Meha Patel, Anna L McNaughton, Jolynne Mokaya, M Azim Ansari, Hizni Salih, Kinga A Várnai, Oliver Freeman, Sarah Cripps, Jane Phillips, Jane Collier, Kerrie Woods, Keith Channon, Jim Davies, Eleanor Barnes, Katie Jeffery, Philippa C Matthews

**Author notes:** These two authors contributed equally (Louise O Downs, David Smith). **CORRESPONDING AUTHOR Email:** **Address:** Medawar Building for Pathogen Research, South Parks Road, Oxford OX1 3SY, UK. **CONFLICT OF INTEREST:** Nil. **FINANCIAL SUPPORT:** LD is funded by the National Institute for Health Research (NIHR). PCM is funded by the Wellcome Trust, grant number 110110. EB is funded by the Medical Research Council UK, the Oxford NIHR Biomedical Research Centre and is an NIHR Senior Investigator. The views expressed in this article are those of the authors and not necessarily those of the NHS, the NIHR, or the Department of Health.

## Abstract

HBsAg and HBeAg have gained traction as biomarkers of control and clearance during chronic hepatitis B virus infection (CHB). Improved understanding of clearance correlates of these proteins could help inform improvements in patient-stratified care and advance insights into underlying mechanisms of disease control, thus underpinning new cure strategies. We collected electronic clinical data via an electronic pipeline supported by the National Institute for Health Research Health Informatics Collaborative (NIHR-HIC), adopting an unbiased approach to generating a robust longitudinal dataset for adults testing HBsAg-positive from a large UK teaching hospital over a six year period (2011-2016 inclusive). From 553 individuals with CHB, longitudinal data were available for 319, representing >107,000 weeks of clinical follow-up. Among these 319 individuals, 13 (4%) cleared HBsAg completely. Among these 13, HBsAg clearance rate was similar in individuals on nucleos(t)ide analogue (NA) therapy (n=4 (31%)), median clearance time 150 weeks) vs those not on NA therapy (n=9 (69%), median clearance time 157 weeks). Those who cleared HBsAg were significantly older, and less likely to be on NA therapy compared to non-clearers (p=0.003 and p=0.001, respectively). Chinese ethnicity was associated with HBeAg positivity (p=0.025). HBeAg clearance occurred both on NA therapy (n=24, median time 49 weeks) and off NA therapy (n=19, median time 52 weeks). Improved insights into the dynamics of these biomarkers can underpin better prognostication and patient-stratified care. Our systematised approach to data collection paves the way for scaling up efforts to harness clinical data to address research questions and underpin improvements in clinical care.

**IMPORTANCE:** Advances in the diagnosis, monitoring and treatment of hepatitis B virus (HBV) infection are urgently required if we are to meet international targets for elimination by the year 2030. Here we demonstrate how routine clinical data can be harnessed through an unbiased electronic pipeline, showcasing the significant potential for amassing large clinical datasets that can help to inform advances in patient care, and provide insights that may help to inform new cure strategies. Our cohort from a large UK hospital includes adults from diverse ethnic groups that have previously been under-represented in the literature. Tracking two protein biomarkers that are used to monitor chronic HBV infection, we provide new insights into the timelines of HBV clearance, both on and off treatment. These results contribute to improvements in individualised clinical care and may provide important clues into the immune events that underpin disease control.

## INTRODUCTION

Chronic HBV (CHB) infection is defined as detectable HBsAg (>20 IU/ml) at ≥2 timepoints ≥6 months apart. Disease activity and treatment response in individuals with CHB infection are most commonly monitored by quantification of HBV DNA viral load (1). However, viral load measurement is expensive, and not universally available, viral DNA levels can fluctuate over time, and quantification can be inaccurate at low levels. Reproducible, automated quantification of other biomarkers such as hepatitis B surface antigen (HBsAg) and/or e-antigen (HBeAg) are therefore attractive biomarkers for use instead of, or alongside, HBV DNA monitoring.

In the context of CHB infection, HBV cccDNA persists as an intranuclear ‘mini-chromosome’ within infected hepatocytes (Fig 1A). HBsAg is produced in excess, from translation of both from the cccDNA reservoir, and from pre-genomic mRNA (2). In a small proportion of cases, HBsAg becomes undetectable over time, suggesting that the cccDNA reservoir is down-regulated, diminished or lost entirely. In the HBV cure field, specific terminology has been adopted to reflect the difference between complete loss of all cccDNA (‘sterilising cure’) vs suppression or dilution of cccDNA to the extent that HBsAg can no longer be detected (‘functional cure’) (3, 4). In practical terms, there is no current way to differentiate between these two outcomes. However, the theoretical distinction is an important one, as sterilising cure reflects complete and permanent loss of HBV from the host, while in the setting of functional cure, there is long-term potential for relapse to occur (best recognised in the setting of immunosuppression (5, 6).

**Fig 1:**
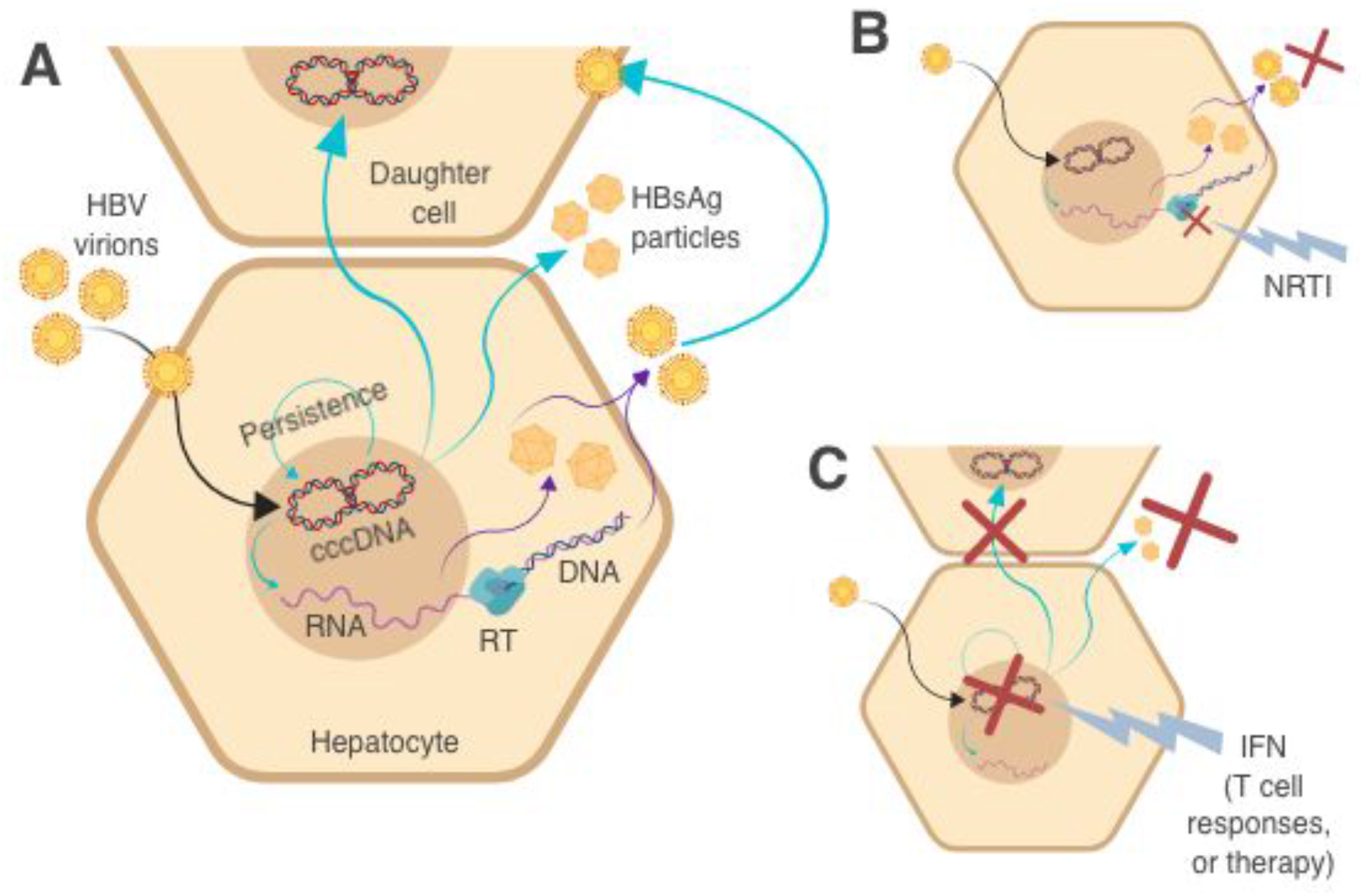
Cartoons depicting key pathways in HBV replication cycle to illustrate targets that may bring about control or clearance. **A: Pathways relevant to maintenance of HBV infection.** HBV viral DNA is released in the nucleus, and cccDNA is formed by covalent ligation of the two DNA strands. A stable minichromosome is formed, allowing persistence of the virus over time. The cccDNA acts as the template for mRNA and pregenomic RNA (pgRNA). Viral reverse transcriptase (RT) generates new genomic DNA from pgRNA. Non-infectious sub-viral particules (SVP) form from HBsAg and new infectious virions assemble, for release into the blood stream. HBsAg measurement accounts for both the SVP and infectious virions, whereas infectious virions alone can be measured through HBV viral load (HBV DNA). **B: Pathways relevant to suppression of HBV infection by NA therapy:** Inhibiton of viral RT suppresses generation of new viral DNA. This means new infectious HBV virions cannot be constructed and HBV DNA is undetectable in plasma. However, cccDNA remains as a persistent reservoir in the hepatocyte nucleus, so HBsAg production can continue and rebound viraemia is likely following cessation of therapy. For this reason, individuals with CHB on successful treatment frequently have an undetectable viral load but remain HBsAg-positive. **C: Pathways relevant to functional or sterilising cure of HBV infection:** Upregulation of host immune responses or therapy with interferon (IFN) leads to elimination of the persistent cccDNA reservoir either through death of the hepatocyte or unknown non-lytic methods. HBsAg and HBV DNA both disappear from the blood stream. In practice, there is no clinical test that can confirm complete (‘sterilising’) cure, so this group is usually regarded as being at a small risk of relapse (i.e. ‘functional’ cure).

Nucleos(t)ide analogues (NA’s) inhibit HBV reverse transcriptase, leading to loss of HBV DNA from the serum, but have no direct effect on cccDNA (3). Thus, HBsAg production can continue unchecked from the cccDNA reservoir, and viral replication frequently returns on cessation of treatment (fig 1B) (3). For this reason, most guidelines currently recommend long term NA treatment (1). Immune responses (either arising naturally or driven by immunotherapy such as interferon) can lead to downregulation or loss of cccDNA to the extent that neither HBsAg nor HBV DNA can be detected in the serum (fig 1C) (4). The long term goal of new immunotherapeutic approaches will be to elicit sterilising cure such that cccDNA is removed with no long-term risk of relapse (7).

HBsAg levels are typically highest in the earlier phases of infection and in HBeAg-positive individuals, frequently correlate with HBV DNA levels in CHB infection, and are associated with risk of subsequent reactivation (8). HBsAg may be a quantifiable risk factor for development of hepatocellular carcinoma (HCC) and chronic liver disease (9), although the relationship is not well defined: in some studies, higher HBsAg levels are associated with lower levels of fibrosis (10–12), while in others, lower baseline HBsAg levels are associated with reduced risk of both cirrhosis and HCC (13). HBsAg levels have also been used to classify individuals into those with inactive carriage (HBV DNA <2000 IU/ml and normal ALT (14, 15)) versus active CHB (with higher viral loads and elevated risks of inflammatory liver disease, fibrosis and cirrhosis (16–19)). HBsAg elimination is widely regarded as a marker of immunological clearance (which may be regarded as ‘functional cure’).

HBeAg-positivity is associated with high viral loads and is therefore a marker of infectivity. Loss of HBeAg is usually associated with production of anti-HBe antibody (a marker of immune-mediated control), and typically associated with lower viral loads. However, although these broad patterns have been described, further efforts are required to elucidate and interpret the dynamics of HBsAg and HBeAg, with the potential to develop insights into the timing and patterns of immunological clearance, and to improve patient-stratified clinical management.

A recent systematic review and meta-analysis has collated literature on HBsAg clearance, with a primary focus on untreated populations (20). This identified 34 studies, but only 14 of these reported ≥2 HBsAg measurements over time, and all but two were in Asia. To ensure we had adequately reviewed the relevant existing evidence on this topic, we also undertook an independent literature review (summarised in Suppl Table 1). We initially identified 43 studies reporting dynamics of HBsAg loss in CHB infection. We excluded studies prior to 2008, those reporting only one HBsAg measurement, and those without an annual or cumulative HBsAg clearance rate, leaving nine relevant studies. As for the meta-analysis, the majority (8/9) were in Asian populations (21–28), with the remaining one based in New Zealand (29). The reported clearance rate of HBsAg ranged from 0.15% per year (27) to 2.7% per year (24) with a maximum cumulative clearance of 3.5% (21). Older age was associated with clearance in two cohorts (23, 29). The role of treatment in clearance is inconsistent, with NA treatment associated with clearance in some cohorts (21, 25) but not in others (26).

HBsAg levels can be used to determine treatment response, although this has been more reliably reported for PEG-IFN2α treatment than for NAs (30, 31), as it implies reduction or removal of the cccDNA reservoir (Fig 1C). Current UK guidelines recommend quantitative HBsAg and HBeAg measurement before starting treatment and at weeks 12, 24 and 48 during treatment, followed by 6 monthly measurement during long term therapy (32). European Association for the Study of the Liver (EASL) guidelines recommend quantitative HBsAg measurement annually in treated patients if HBV DNA is undetectable, as well as using HBsAg levels to inform the decision to stop treatment (1). EASL guidelines also recommend HBeAg measurement as part of the initial clinical assessment, and list HBeAg loss as one of the serological responses to treatment, but do not specify a frequency for follow-up testing (1).

International targets arising from the United Nations ‘sustainable development goals’ have set a challenge for elimination of CHB infection as a public health threat by the year 2030 (33). Recognising the multi-lateral approaches that will be required to reach this ambitious goal, we here focus on two inter-related aims:

i. We set out to showcase how longitudinal data for individuals with CHB can be collected through an unbiased electronic pipeline that collates, cleans and anonymises routinely-collected electronic clinical data, in this case driven by infrastructure supported by the UK National Institute for Health Research (NIHR) Health Informatics Collaborative (HIC); (www.hic.nihr.ac.uk). The aim is to harness clinical data to drive research and quality improvements in diagnostics, monitoring and therapy of viral hepatitis, and to underpin new questions for basic science. Through developing and testing this system, we have devised an approach that can be rolled-out to incorporate other centres, with substantial gains predicted through the power of large datasets.
ii. We analysed data for HBV sourced from a tertiary referral UK teaching hospital, in order to develop better insights into patterns of HBsAg and HBeAg clearance. Through the application of an unbiased approach (agnostic to treatment, clinical stage of disease, other biomarkers, or genotype of infection), we aim to develop a clear picture of the dynamics of clearance. Identifying demographic or clinical characteristics that predict specific disease outcomes, provides opportunity for the investigation of immunological correlates of control and clearance.

Collectively, this enterprise provides proof-of-principle for the systematic use of electronic clinical data in informing studies of viral hepatitis, as well as shedding new light on the dynamics of clearance of HBsAg and HBeAg.

## RESULTS

### Description of a clinical cohort of chronic HBV infection

We identified 553 individuals who tested HBsAg-positive during the six-year period 2011-2016, inclusive. Of these, 319 met inclusion criteria for further analysis (as shown in Table 1; Fig 2). Characteristics of the cohort are summarised in Table 2 and the complete metadata for these 319 CHB patients is available as a supporting data file (Suppl Table 2). We collected longitudinal data for a total of 107,702 person-weeks (range 61-702 weeks, mean 338 weeks (6.5 years) of follow-up per individual, IQR 174-487). The median age at first HBsAg test was 34 years (IQR 29 – 43, range 10 – 71), and males accounted for 191/319 (60%) of cases. HIV co-infection was documented in 9 individuals (2.8%), although we cannot exclude the possibility that the true prevalence of HIV-coinfection was higher due to a proportion of individuals who did not have a recent HIV test result.

**Fig 2:**
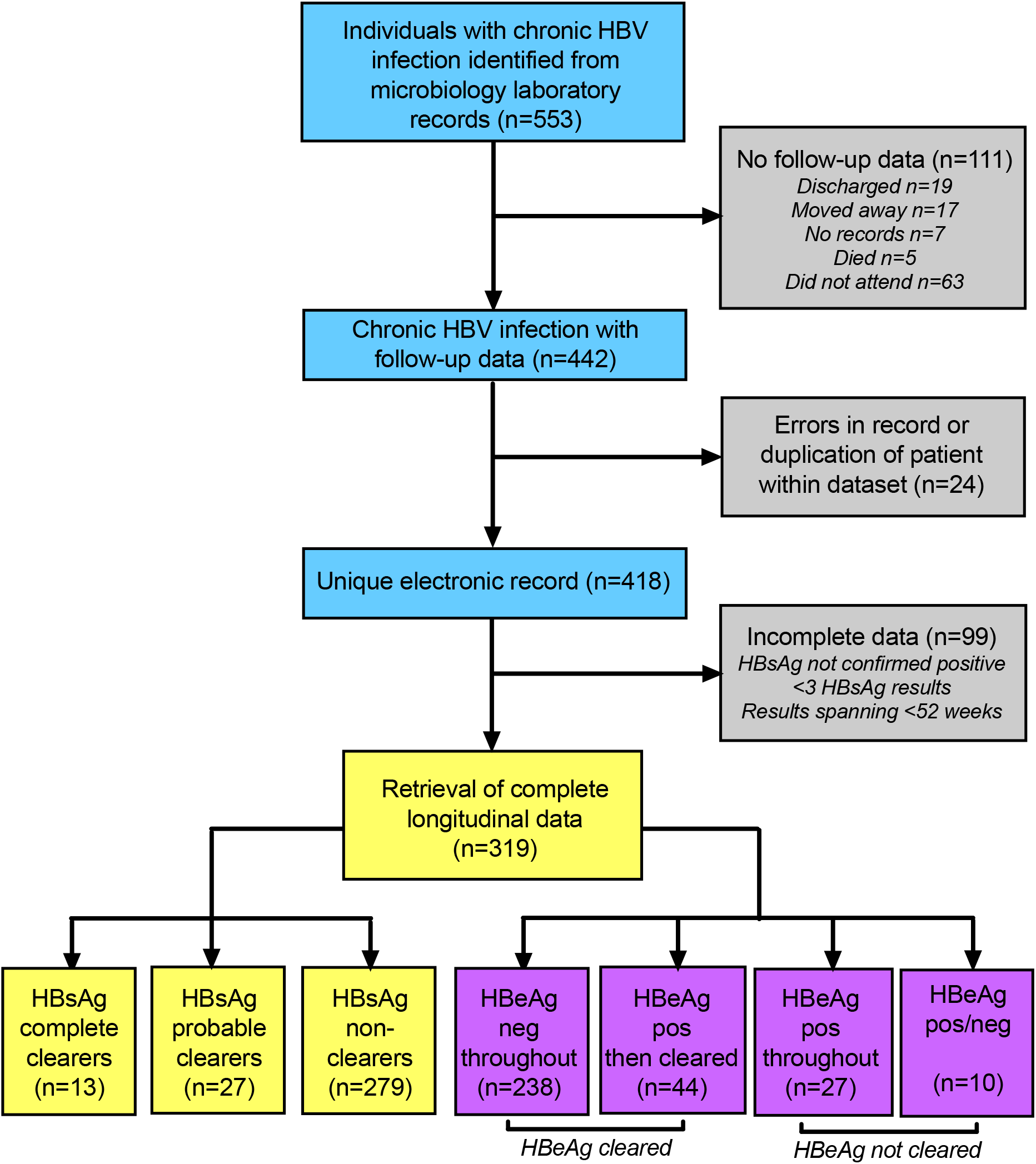
Flowchart showing identification and classification of adults with chronic HBV infection from a hospital electronic system. The figure represents 319 individuals who met inclusion criteria, and divides these into three different categories according to HBsAg clearance, and four categories for HBeAg; (for classification criteria, see Table 1).

**Table 1:**
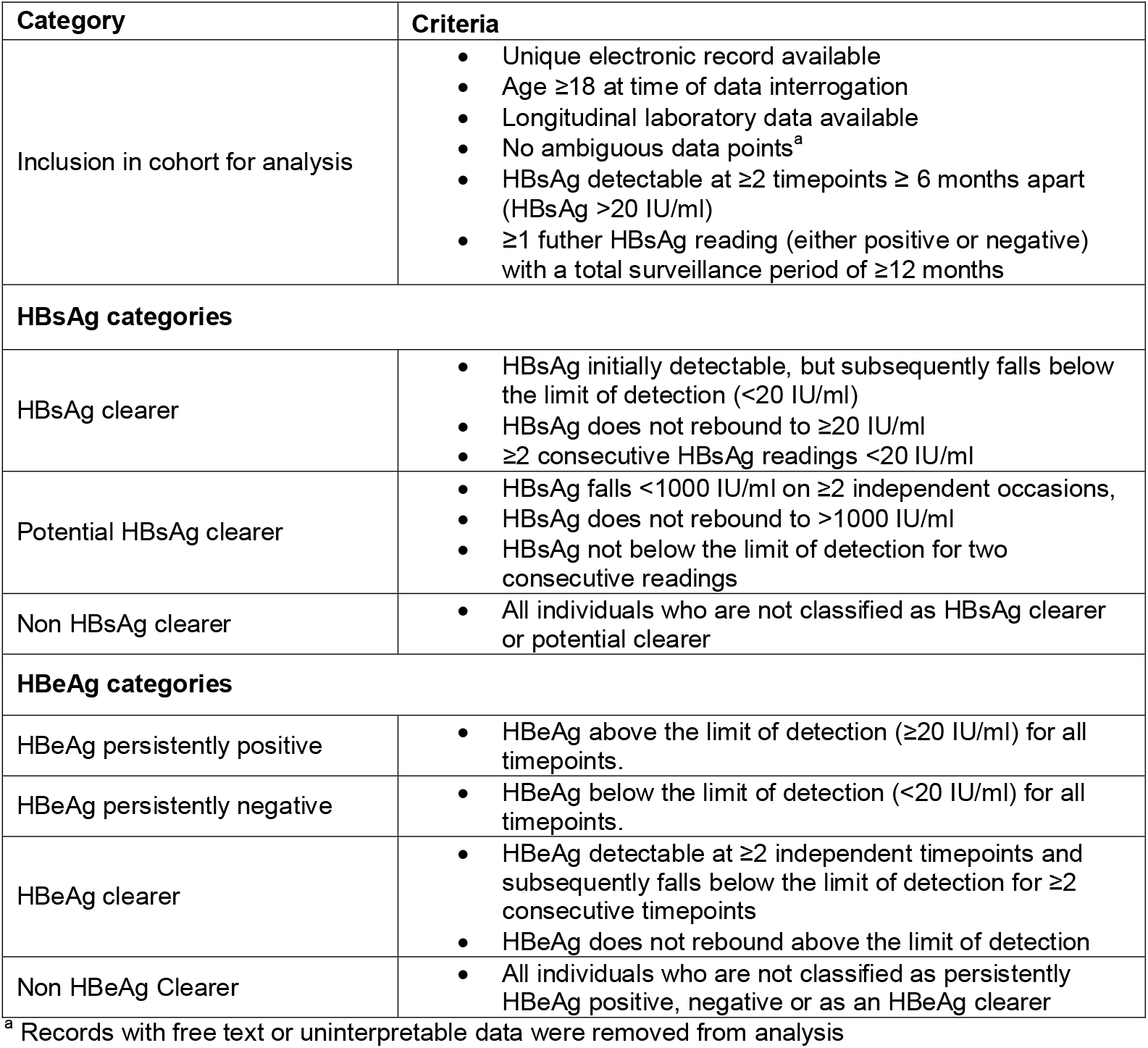
Summary of criteria used to confirm inclusion in the analysis and to classify individuals according to HBsAg dynamics and HBeAg dynamics

| Category | Criteria |
| --- | --- |
| Inclusion in cohort for analysis | • Unique electronic record available<br>• Age $\geq 18$ at time of data interrogation<br>• Longitudinal laboratory data available<br>• No ambiguous data points<sup>a</sup><br>• HBsAg detectable at $\geq 2$ timepoints $\geq 6$ months apart (HBsAg $> 20$ IU/ml)<br>• $\geq 1$ further HBsAg reading (either positive or negative) with a total surveillance period of $\geq 12$ months |
| **HBsAg categories** |  |
| HBsAg clearer | • HBsAg initially detectable, but subsequently falls below the limit of detection ($< 20$ IU/ml)<br>• HBsAg does not rebound to $\geq 20$ IU/ml<br>• $\geq 2$ consecutive HBsAg readings $< 20$ IU/ml |
| Potential HBsAg clearer | • HBsAg falls $< 1000$ IU/ml on $\geq 2$ independent occasions,<br>• HBsAg does not rebound to $> 1000$ IU/ml<br>• HBsAg not below the limit of detection for two consecutive readings |
| Non HBsAg clearer | • All individuals who are not classified as HBsAg clearer or potential clearer |
| **HBeAg categories** |  |
| HBeAg persistently positive | • HBeAg above the limit of detection ($\geq 20$ IU/ml) for all timepoints. |
| HBeAg persistently negative | • HBeAg below the limit of detection ($< 20$ IU/ml) for all timepoints. |
| HBeAg clearer | • HBeAg detectable at $\geq 2$ independent timepoints and subsequently falls below the limit of detection for $\geq 2$ consecutive timepoints<br>• HBeAg does not rebound above the limit of detection |
| Non HBeAg Clearer | • All individuals who are not classified as persistently HBeAg positive, negative or as an HBeAg clearer |

<sup>a</sup> Records with free text or uninterpretable data were removed from analysis

**Table 2:** Baseline characteristics of 319 individuals with chronic HBV infection recruited through a UK cohort and classified according to pattern of HBsAg clearance over time.

|  | Whole cohort | HBsAg clearers and potential clearers | HBsAg non-clearers | p-value (uni-variable analysis) | p-value (multi-variable analysis) | Effect Size | Std Error |
| --- | --- | --- | --- | --- | --- | --- | --- |
| Number of individuals | 319 | 40 | 279 | NA | NA |  |  |
| Median age in years at time of first HBsAg test | 34 | 40 | 34 | 0.0034* | 0.0081* | -0.4 | 0.01 |
| Sex (%) |  |  |  | 0.605 | 0.605 | 0.14 | 0.36 |
| Male | 191 (60) | 26 (65) | 165 (59) |  |  |  |  |
| Female | 128 (40) | 14 (35) | 114 (41) |  |  |  |  |
| Self-reported ethnicity (%) |  |  |  |  |  |  |  |
| White | 92 (29) | 15 (38) | 77 (28) | 0.649 | 0.632 | -0.55 | 0.55 |
| Mixed | 18 (6) | 0 (0) | 18 (6) | 0.991 | 0.989 | 15.39 | 898.69 |
| Asian or Asian British | 52 (16) | 7 (18) | 45 (16) | 0.641 | 0.705 | -0.25 | 0.63 |
| Black or Black British | 46 (14) | 5 (12) | 41 (15) | 0.697 | 0.638 | -0.06 | 0.68 |
| Chinese | 56 (18) | 8 (20) | 48 (17) | 0.902 | 0.914 | -0.28 | 0.62 |
| Any Other Ethnic Group | 7 (2) | 0 (0) | 7 (3) | 0.992 | 0.994 | 15.39 | 1458.87 |
| Not Stated | 48 (15) | 5 (12) | 43 (15) | NA | NA | NA | NA |
| HBeAg positive status at baseline (%) | 81 (25) | 6 (15) | 65 (23) | 0.3105 | 0.131 | 0.51 | 0.49 |
| Median elastography score, kPa (most recent value) | 5.3 | 4.5 | 5.5<sup>a</sup> | 0.18 | NA | NA | NA |
| Number of patients receiving treatment (%) | 142/211<sup>b</sup> (67) | 11/40<sup>c</sup> (28) | 131/171<sup>b</sup> (76) | NA | NA | NA | NA |

<sup>a</sup> Elastography data available for 42 individuals in the non-clearance group, as data not routinely recorded electronically.
<sup>b</sup> Treatment data were missing for 108 individuals among the HBsAg non-clearers, as data not routinely recorded electronically.
<sup>c</sup> Treatment in the 12 months before the last positive HBsAg test
NA = not applicable

### Frequency of HBsAg clearance

Exemplar patterns of HBsAg clearance are illustrated in Fig 3 (as per definitions in Table 1). Using the most stringent definition of HBsAg clearance, we documented complete clearance in 13/319 (4.1%) individuals (for full details see Suppl Table 2 and clearance trajectories shown in Suppl Fig 1). The HBsAg clearance rate for this cohort was 0.6% per year. In only 2/13 cases could we estimate the likely duration of infection prior to clearance, one individual who had been vertically infected (HBS-145) and one with iatrogenic infection related to a blood transfusion in childhood (HBS-113). These individuals were both infected for approximately 25 years before clearing HBsAg.

**Fig 3:**
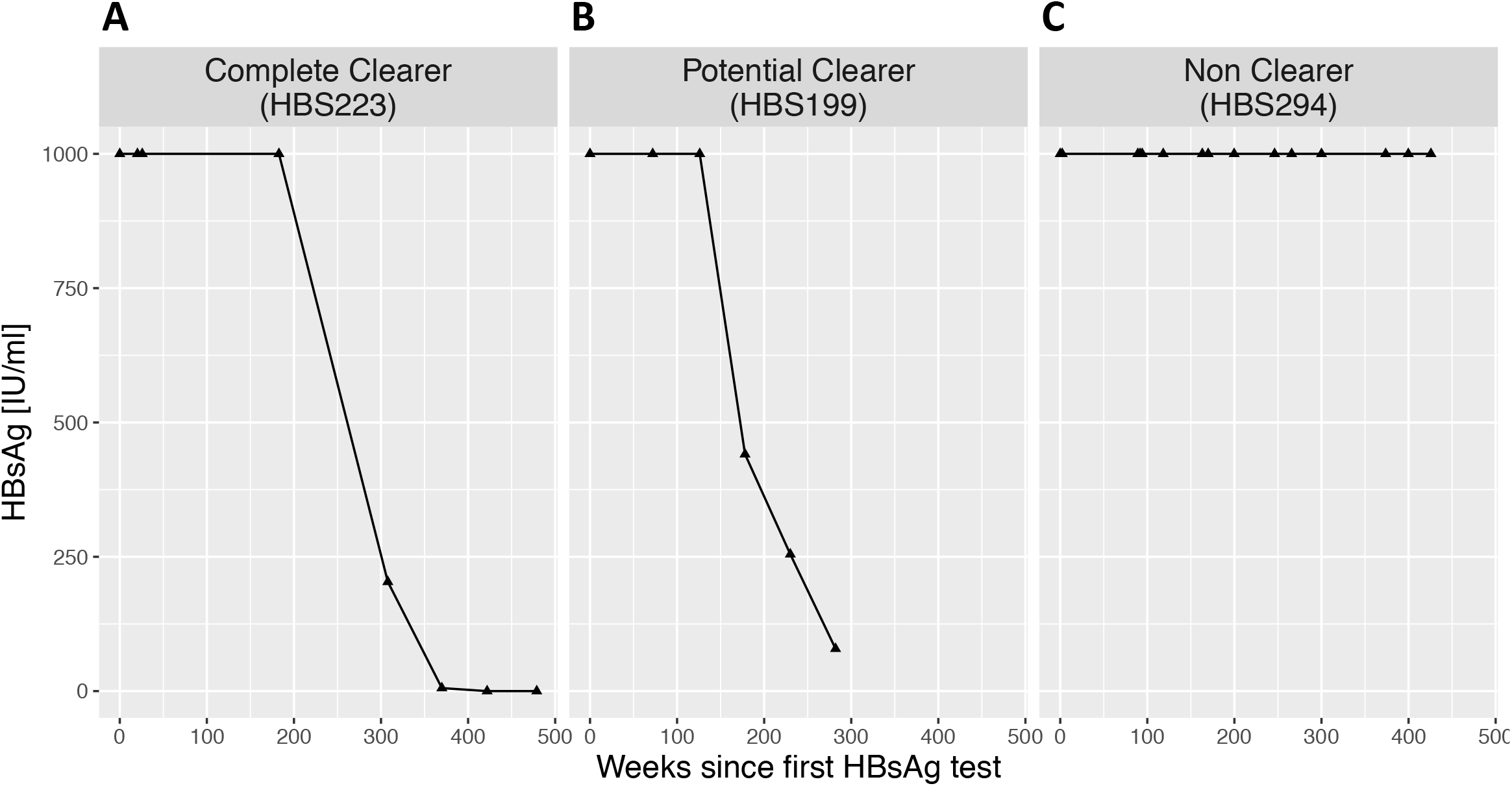
Exemplar trajectories of HBsAg over time representing adults with chronic HBV infection. Individuals are classified as (A) a complete HBsAg clearer, (B) a potential HBsAg clearer (C) a non-HBsAg clearer; (for classification criteria, see Table 1).

We classified an additional 27/319 (8.5%) individuals as ‘potential clearers’ on the grounds of HBsAg trends consistently declining towards clearance (criteria in Table 1; clearance curves shown in Suppl Fig 2). These represent a more heterogenous group, but the clearance trajectory in all cases suggests that they would meet the more stringent clearance criteria if prospective surveillance were to be continued. In contrast, HBsAg curves for non-clearers are shown in Suppl Fig 3.

### Characteristics of individuals with HBsAg clearance or potential clearance

Adults classified as completely or potentially clearing HBsAg were significantly older than non-clearers (median age 40 vs 34 years; p=0.003; Table 2; Suppl Fig 4). There was no difference in sex or ethnic origin between individuals in different HBsAg clearance categories (Table 2). The majority of those who completely cleared HBsAg were HBeAg-negative throughout the period of observation (10/13, 77%). Among the remaining three with detectable HBeAg, two of these lost HBeAg prior to clearing HBsAg (HBS-197 and HBS-223), while one (HBS-195) cleared HBsAg and HBeAg together (Suppl Fig 1). In three cases (HBS-113, HBS-145 and HBS-195), HBV DNA was cleared at the same time as HBsAg; however, in the other ten individuals (77% of clearers), HBV DNA levels were low (<100 IU/ml) throughout the period of HBsAg clearance.

### Rate of HBsAg clearance

HBsAg clearance occurred over a median time of 157 weeks (95% CI 90-239 weeks) (Fig 4A). Comparing individuals on treatment (n=4) vs. off treatment (n=9) during or in the 12 months prior to HBsAg clearance, clearance occurred over similar time frames (median 150 weeks in those on treatment vs. 157 weeks in those not on treatment; Fig 4B). Among 279 HBsAg ‘non-clearers’, 246/279 (88%) had HBsAg levels that were persistently >1000 IU/ml. The remaining 12% had more heterogenous HBsAg dynamics, including transient dips <1000 IU/ml (e.g. HBS-298) and sustained levels <1000 IU/ml but without a trend towards clearance (e.g. HBS-368).

**Fig 4:**
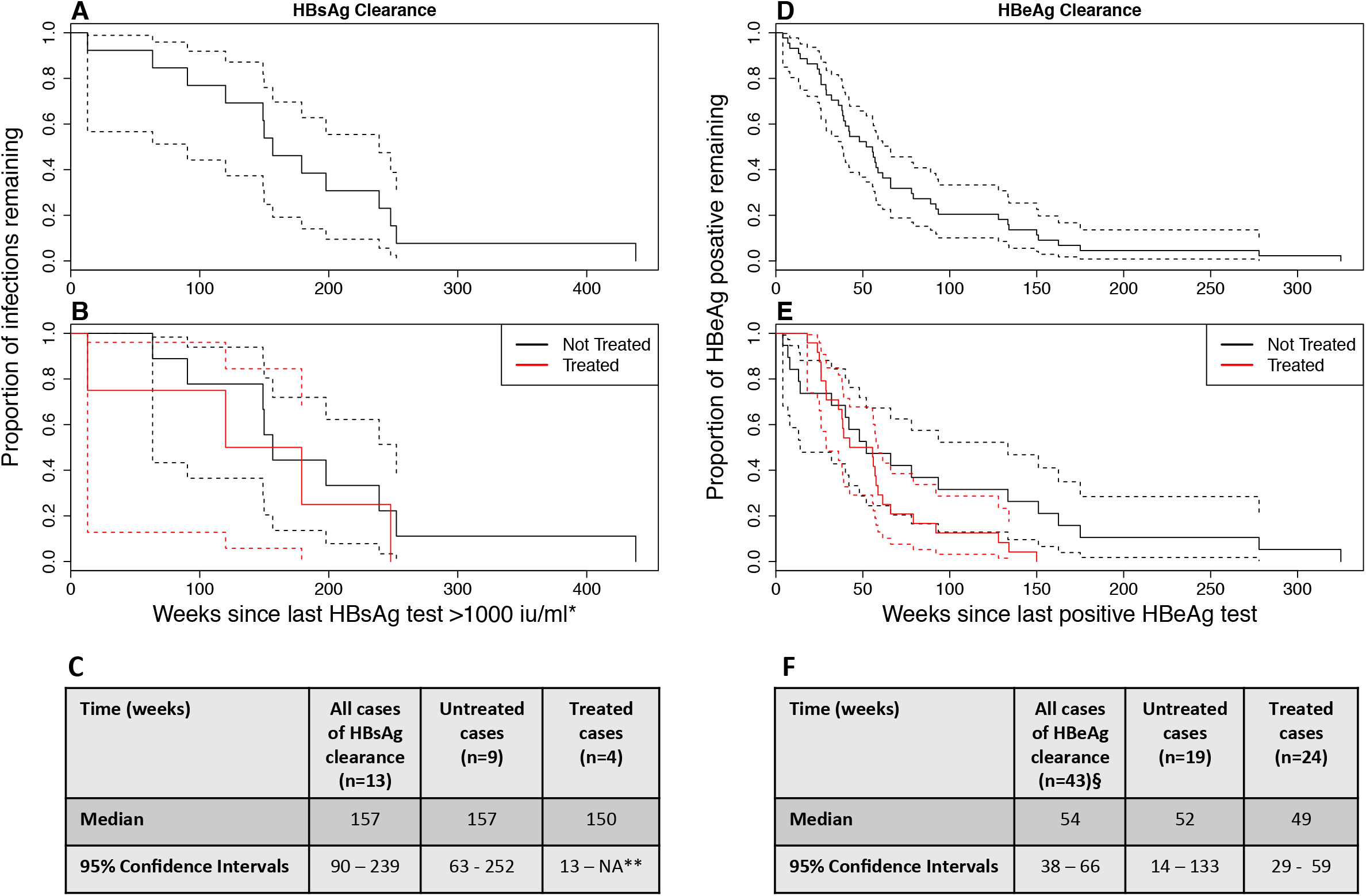
Kaplan-Meier curves showing trajectory of HBsAg clearance (N=13) and HBeAg clearance (N=43) for selected individuals who met criteria for complete clearance from within a cohort of adults with chronic HBV infection. Data are shown for HBsAg (panels A-C) and for HBeAg (panels D-F), initially for all clearers (panels A and D), and then subdivided according to treatment status (panels B and E). Boxes C and F report the median time to clearance for each group in weeks, with 95% confidence intervals. For HBsAg clearance, the upper confidence interval for treated cases cannot be determined due to small numbers. Treatment of HBsAg clearers and potential clearers comprised TDF monotherapy (n=3), TDF with emtricitabine (n=2), 3TC with ADV or TDF (n=4), 3TC monotherapy (n=1), ETV montherapy (n=1). Treatment of HBeAg clearers comprised TDF monotherapy (n=10). 3TC monotherapy (n=2) ETV monotherapy (n=5), 3TC with ADV (n=3), IFN with RBV (n=1), IFN monotherapy (n=3), treatment data were not available for one individual. * When no values >1000 IU/ml were recorded, the highest value was used. ** Not enough data to cacluate upper CI. § Treatment status not known for 1 individual.

### Treatment status of HBsAg clearers vs non-clearers

During the HBsAg clearance phase, or in the 12 months prior, 4/13 (31%) individuals defined as having completely cleared HBsAg were on NA therapy (Fig 4B,C). These individuals had received treatment for a median of 13 months (range 2 months – 8 years) prior to clearance. The other nine (69%) were not on treatment in the 12 months prior to HBsAg clearance, but one had received PEG-IFNα therapy 4 years earlier. In those individuals defined as ‘potential clearers’, 7/27 (26%) received NA treatment, two of whom received TDF as part of an HIV treatment regimen.

We also reviewed treatment data for the 279 individuals who did not clear HBsAg, and were able to retrieve data for 171 of these (61%). Among these, 131 (77%) had received treatment of some type, and 40 had never been treated (23%). We were not able to determine robust time-frames for most treatment episodes. Based on these data, non-HBsAg clearers were statistically more likely to be on treatment than HBsAg clearers (131/171, vs 4/13 respectively, p=0.001 by Fisher’s Exact test). This may reflect inherently better immune control in the group who clear HBsAg, meaning they are less likely to meet criteria for treatment than non-clearers. However, these data must be interpreted with caution, as bias is introduced as a result of missing data among the non-clearers, and by different time-lines for follow-up (we assessed treatment cross-sectionally in clearers based on a specific time of HBsAg loss, for which there is no equivalent among non-clearers, thus we may have assessed longer follow-up times in the latter group).

### HBeAg status

HBeAg was detectable in 81/319 (25%) of individuals at the start of the observed time period. Among these, 51/81 (63%) were male and the median age was 34. By multivariable analysis, Chinese ethnicity was associated with HBeAg-positive status, with 22/56 (39%) of Chinese individuals being HBeAg-positive (p=0.025). We documented HBeAg clearance in 44/81 (54%) of these individuals over the observed time period (Table 3) at a median age of 37 years. HBeAg loss occurred over a median period of 54 weeks (95% CI 38-66 weeks) between last positive and first negative HBeAg test (Fig 4D). Median clearance was 49 weeks (95% CI 29-59 weeks) for individuals who had received treatment in the year prior to the last positive HBeAg result (n= 24, 55%) and 52 weeks (95% CI 14-133) for untreated individuals (n=19, 43%); treatment data were not available for 1 individual (Fig 4E,F). We also reviewed treatment data for those who did not clear HBeAg, and were able to retrieve data for 27 of these (73%). Of these, 24 (89%) had received some treatment whilst 3 (11%) were untreated.

**Table 3:** Baseline characteristics of HBeAg positive individuals classified according to HBeAg clearance over the observed time period

|  | HBeAg clearers | HBeAg non-clearers | p-value (uni-variable analysis) |
| --- | --- | --- | --- |
| Number of individuals | 44 | 37 |  |
| Median age in years at time of first HBsAg test | 34 | 35 | 0.75 |
| Sex (%) |  |  | 1 |
| Male | 29 (66) | 25 (58) |  |
| Female | 15 (34) | 12 (32) |  |
| Self-reported Ethnicity (%) |  |  |  |
| White | 12 (27) | 10 (27) | 0.959 |
| Mixed | 4 (9) | 4 (11) | 0.819 |
| Asian or Asian British | 8 (18) | 3 (8) | 0.427 |
| Black or Black British | 6 (14) | 0 (0) | 0.991 |
| Chinese | 8 (18) | 14 (38) | 0.330 |
| Any Other Ethnic Group | 1 (2) | 2 (5) | 0.512 |
| Not Stated | 5 (11) | 4 (11) | NA |
| Median elastography score, kPa (based on most recent value) | 5.5 | 4.55 | 0.24 |
| Number of patients receiving treatment <sup>a</sup> (%) | 24/44 (55) | 24/27<sup>b</sup> (89) | NA |

NA = not applicable
<sup>a</sup> Treatment in the 12 months prior to the last positive HBeAg result
<sup>b</sup> Treatment data were missing for 10 individuals among the HBeAg non-clearers as data not routinely collected electronically.

### Association between HBsAg clearance and ALT

Complete longitudinal ALT data are shown for each individual in Suppl Figs 1–3. We investigated whether there were differences in ALT according to HBsAg clearance (for each of the three HBsAg groups defined in Fig 2). There was no significant difference in ALT at the time of first test between HBsAg clearers, ‘potential clearers’ and non-clearers (data not shown). ALT data were available before and during HBsAg clearance for 11/13 individuals who cleared HBsAg. Among these, three individuals (HBS-162, HBS-195 and HBS-314) had a spike in ALT before clearance which returned to the normal range after HBsAg clearance. Another individual (HBS-230) also had a slightly raised ALT before HBsAg clearance, but this did not normalise after HBsAg clearance. In the 7 other cases, ALT results remained within the local reference range (10-45 iu / L) for the entire period of surveillance (Suppl Fig 1).

### Relationship between HBsAg and HBV DNA

In 11/13 HBsAg clearers, HBV DNA was below the limit of detection (<20 IU/ml) throughout; in two cases, HBV DNA was cleared at the same time as HBsAg (Suppl Fig 1). The HBV DNA trajectory of individuals classified as potential clearers was more heterogenous (Suppl Fig 2): 10 individuals had cleared HBV DNA by the time of their last HBsAg test, 9 had negative HBV DNA results at some point but had subsequent detectable viraemia, and 8 individuals had detectable HBV DNA throughout the period of surveillance.

## DISCUSSION

### Novelty

HBsAg clearance in CHB is an uncommon event, and large cohorts over a long period of clinical follow up are therefore required to describe the characteristics of individuals who clear, and to determine the specific dynamics of serological changes. Although there have been previous studies reporting HBsAg loss, our literature review confirms that these are mostly focussed in Asia, and that relatively few studies track longitudinal data in an unbiased way. Our current approach adds novelty in a variety of ways:

i. We apply a new bespoke, algorithmic approach to collating a large longitudinal clinical dataset from multiple electronic sources. This allowed us to make use of data that are generated by routine clinical laboratories, but are not routinely used for patient care such as quantiative measures of HBsAg and HBeAg. This method of data collection also facilitated the robust identification and exclusion of duplicate patient records.
ii. Undertaking this analysis in a UK-based cohort provides a novel and more diverse mixture of host ethnicities (and by inference, diverse viral genotypes). To our knowledge this is one of the only studies of this kind in such a population.
iii. We report ≥2 HBsAg timepoints for each individual, providing long periods of clinical follow-up and the opportunity to track uncommon clearance events over time;
iv. Unlike some previous studies of HBsAg clearance that introduce bias through a focus on treatment or based on patient recall for follow-up, the approach we took is agnostic to other parameters, thereby providing a more inclusive picture of all individuals with CHB infection;
v. In addition to reporting longitudinal data for HBsAg loss, we also track HBeAg loss over time. HBeAg loss is an important immunological event (34) signifying control (typically in association with a fall of HBV DNA levels), and may also be an important target for interventions at a population level (35).

### The value of the NIHR HIC approach

The NIHR HIC approach, involving the generation of standardised datasets based upon routinely-collected data, focussed on the needs of researchers in particular clinical and/or therapeutic areas, supports the re-use of tools, data, and insights across multiple research projects and organisations. As we address a wider range of questions, and as we continue to share expertise with other university-hospital partnerships, we will increase the breadth, depth, and quality of the dataset, covering a wider range of variables, for a larger patient population, and recording more information about the provenance and interpretation of the values obtained. All of the data used for this paper, and our understanding of that data, will be available for use by other researchers. Any questions that we were unable to fully address, and other questions that emerged during the analysis, will help to inform the future development of the dataset.

### Role of treatment in HBsAg clearance

Our dataset corroborates prior literature in confirming that treatment is not pre-requisite for clearance, and that immunological clearance of HBsAg and HBeAg can occur independently of antiviral therapy (Fig 1C) (23, 36). There are multiple host and viral factors influencing outcome during CHB infection including host factors such as age, obesity, gender and diabetes along with genetic variations in CD8+ T-cell responses (mediated by HLA genotype), T-cell receptor antagonism and viral escape mutations (37, 38).

Due to small numbers, we did not have statistical power to determine whether there was a significant difference in the time taken to clear either HBsAg or HBeAg in individuals on treatment compared to an untreated group. However, the comparable speed of clearance on vs off treatment suggests that clearance trajectories are similar irrespective of NA treatment. We found that NA treatment was more common among non-clearers, which may reflect a genuinely higher proportion of this group meeting treatment criteria, but may also be biased by the incomplete nature of our treatment data. Further prospective studies are needed to study the relationship between clearance and treatment in more detail.

### Timing of HBsAg and HBeAg clearance

Based on the epidemiology of HBV infection in this cohort, in which a substantial proportion of individuals are likely to have been infected at birth or in early childhood, it is intriguing that HBsAg and HBeAg clearance occur apparently at random in middle adulthood. In the case of HBsAg, clearance, its association with older age has been previously reported in studies we identified through our literature review (20, 23, 29). The chances of clearance may be cumulative over time; individuals infected for longer periods of time could thus have a higher chance of clearance, explaining why individuals who clear infection are, on average, older than non-clearers.

HBeAg clearance occurred over a median period of 54 weeks, substantially more quickly than HBsAg clearance which was documented over a median period of 157 weeks, perhaps indicating different underlying mechanisms at play (34, 37, 39). Further studies are needed to determine the relevant immune responses that underpin this clearance, and to identify possible triggers for clearance.

### Relevance of HBsAg and HBeAg for clinical practice and research

While some guidelines recommend monitoring of HBsAg levels (1, 40), there is a lack of consistent understanding about how to interpret individual or longitudinal measurements. Developing better insights into the prognostic information that can be captured from this biomarker could be relevant to predicting patient outcomes and providing stratification of therapy. In this study, we did not have routine access to HBsAg levels >1000 IU/ml, but as these data progressively become available, future studies will have the opportunity to develop a better picture of HBsAg distribution across the whole range of CHB infections. Advocacy is required to provide more universal access to platforms that quantify HBsAg, and to improve clinical practice through interval measurements of HBsAg in chronically infected patients.

The picture we have developed here suggests that the majority of individuals who develop a sustained pattern of HBsAg decline below 1000 IU/ml are likely to go on to clear HBsAg, consistent with previous longitudinal surveillance suggesting that baseline HBsAg levels may be a more accurate prognostic marker than HBV viral load (28, 29). Prospective studies of large HBV cohorts are likely to be needed to identify individuals on a clearance trajectory; enhanced surveillance of these individuals is a promising future route to understanding the immunological correlates of HBsAg clearance.

### Caveats and limitations

Routinely-collected clinical data may be lacking in context, consistency, and completeness. Health professionals recording information to support decision-making and continuity of care, and the systems that they use, may fail to record additional, contextual information needed to address specific research questions. Variations in practice may mean that data from different sites, or from different clinicians, are incompatible (see Table 4). A collaborative approach to data quality improvement, with substantial, local clinical engagement, will help to address these challenges, but there is always more to be done, not least, as practice changes and new research questions evolve. For this paper, the questions that we were able to ask, and the size of the population considered, were limited by the nature and means of the data recorded, rather than by the basic availability. We considered only those clinical records for which the data were sufficiently complete, and for which the context was adequately explained, accepting the possibility that our exclusion of other records could introduce systematic bias.

**Table 4:** Factors influencing the analysis of retrospective clinical HBV data

| Category of influence | Examples of the effect on data integrity |
| --- | --- |
| **Patient factors** | • Many individuals with CHB infection globally are not diagnosed; those with data available for clinical analysis represent a distinct minority group who have been able to access healthcare and follow-up (44).<br>• Patients are lost to follow-up or move between regions.<br>• HBV diagnosis rarely occurs in acute infection, so the duration of infection prior to clearance is unknown.<br>• HBsAg clearance is a relatively infrequent event and thus patient numbers for analysis are small.<br>• Description of a changing cohort is challenging e.g. age changes over time, patients start and stop therapy. |
| **Healthcare factors** | • Different assays are not always requested simultaneously, thus limiting the correlation between variables (e.g. HBV DNA vs HBsAg).<br>• Follow-up occurs over a variety of different time frames, with different intervals between follow up visits; clearance durations may therefore be over-estimated due to infrequent sampling.<br>• Treatment can alter the dynamics of biomarkers (e.g. ALT, HBV DNA). |
| **Laboratory factors** | • Assay platforms change over time, which may alter sensitivity, specificity and limits of detection.<br>• Quantitative assays have upper and lower limits of quantification; values outside the window of detection cannot be analysed.<br>• False positive or false negative tests may occur.<br>• Certain data are not routinely generated or captured (e.g. HBV genotype). |
| **Data factors** | • Results are captured by a variety of different electronic systems (electronic patient record, electronic laboratory systems, pharmacy systems, hand-written clinical notes, dictated clinic letters).<br>• Different healthcare professionals may not record data consistently and coding is subject to errors.<br>• Free text entries in laboratory reporting can lead to errors or ambiguities (e.g. use of comma vs. full stop for decimal point). Certain parameters are not consistently recorded, e.g. ethnicity.<br>• The electronic pipeline only collects certain pre-defined data (e.g. for HIV, HCV, HDV we were only able to access viral load data, not antibody tests, and therefore we do not know the denominator of total tests performed).<br>• Treatment data may not be recorded electronically (often recorded as part of paper notes, making them more difficult to trace); start dates often not documented for patients on long-term treatment.<br>• Poor continuity of data when patients are transferred between different healthcare providers. |

### Future questions

Prospective surveillance is important in order to provide the opportunity for studying relevant immune responses during the clearance phase. As we have shown that clearance is a relatively long process, occurring over a median of 54 weeks for HBeAg and 157 weeks for HBsAg, this provides a window of opportunity for sampling and follow-up. There is an important distinction to be made between functional cure (sustained loss of HBsAg) and sterilising cure (loss of cccDNA integrated into hepatocyte nuclei), and interest in how to determine these different outcomes. There are currently many new therapeutics in clinical trials aimed at targeting cccDNA directly including capsid effectors, RNA interference and gene editing (7). Further work is needed to develop biomarkers that can detect cccDNA in order to distinguish between these two different outcomes.

Studies of both host and viral genetics are required to underpin a better understanding of the mechanisms of clearance, including new approaches to generating full length deep sequencing of HBV, and unbiased methods to study host genetic polymorphisms that impact on disease outcome. In order to power such studies sufficiently to detect relevant signals, large collaborative multi-centre studies may be required. As we improve our insights into the dynamic changes of serological markers, opportunities arise for improving prognostication and providing better patient-stratified care.

## MATERIALS AND METHODS

### Health Informatics Collaborative Infrastructure

The UK National Institute for Health Research (NIHR) Health Informatics Collaborative (HIC) (www.hic.nihr.ac.uk) is a programme of infrastructure development aimed at increasing the quality and availability of routinely-collected clinical data for translational research. Eighteen university-hospital partnerships across England have signed a framework data sharing agreement, and are working to facilitate the sharing and re-use of data across centres, for approved research purposes. A key component of the NIHR HIC approach is the creation of standardised datasets to support research in specific therapeutic areas, with relevant variations in context and practice recorded as structured metadata, facilitating re-use at scale. Viral hepatitis was selected as one of the initial areas for infrastructure development: in Oxford, this has led to the establishment of new data flows from clinical and laboratory systems, the design of new screens for data capture, and the implementation of several, programmatic (and re-usable) data transformations.

### Clinical cohorts

Our HBV cohort was collected from the records of a large UK teaching hospital in Oxford (http://www.ouh.nhs.uk/), which provides 1 million patient contacts per year and receives laboratory samples from the community and four inpatient sites. We retrospectively identified individuals aged ≥18 at time of database interrogation (26-Mar-2018) with chronic HBV infection (defined as positive HBsAg on ≥2 occasions ≥6 months apart) based on laboratory data collected between January 2011 and December 2016. Inclusion criteria and other case definitions are set out in Table 1. It is standard practice in Oxfordshire to start patients on HBV treatment based on UK NICE guidelines which determine treatment eligibility using age, sex, ALT, HBV DNA and fibroscan score (40).

### Data collection

Our cohort was initially defined by an electronic search of the microbiology laboratory information management systems (LIMS) to identify individuals with a positive HBsAg test. Individual subjects were allocated a pseudo-anonymised ID prefixed ‘HBS’, these ID numbers are included in the text to allow relevant results to be identified from within our metadata table (Suppl data table 2). We generated a data specification using search terms (Table 5) to define the data set. The Oxford NIHR Biomedical Research Centre (BRC) data warehouse receives data from operational systems within the hospital, such as electronic patient records (EPR) and LIMS (Fig 5). Within the warehouse, the data are linked, transformed, and reorganised to better support the generation of data products focussed upon a particular purpose or research area. In this case, the data product is a database containing de-identified information on patients with hepatitis. These data were cleaned and individuals not meeting inclusion criteria (Table 1) were removed.

**Fig 5:**
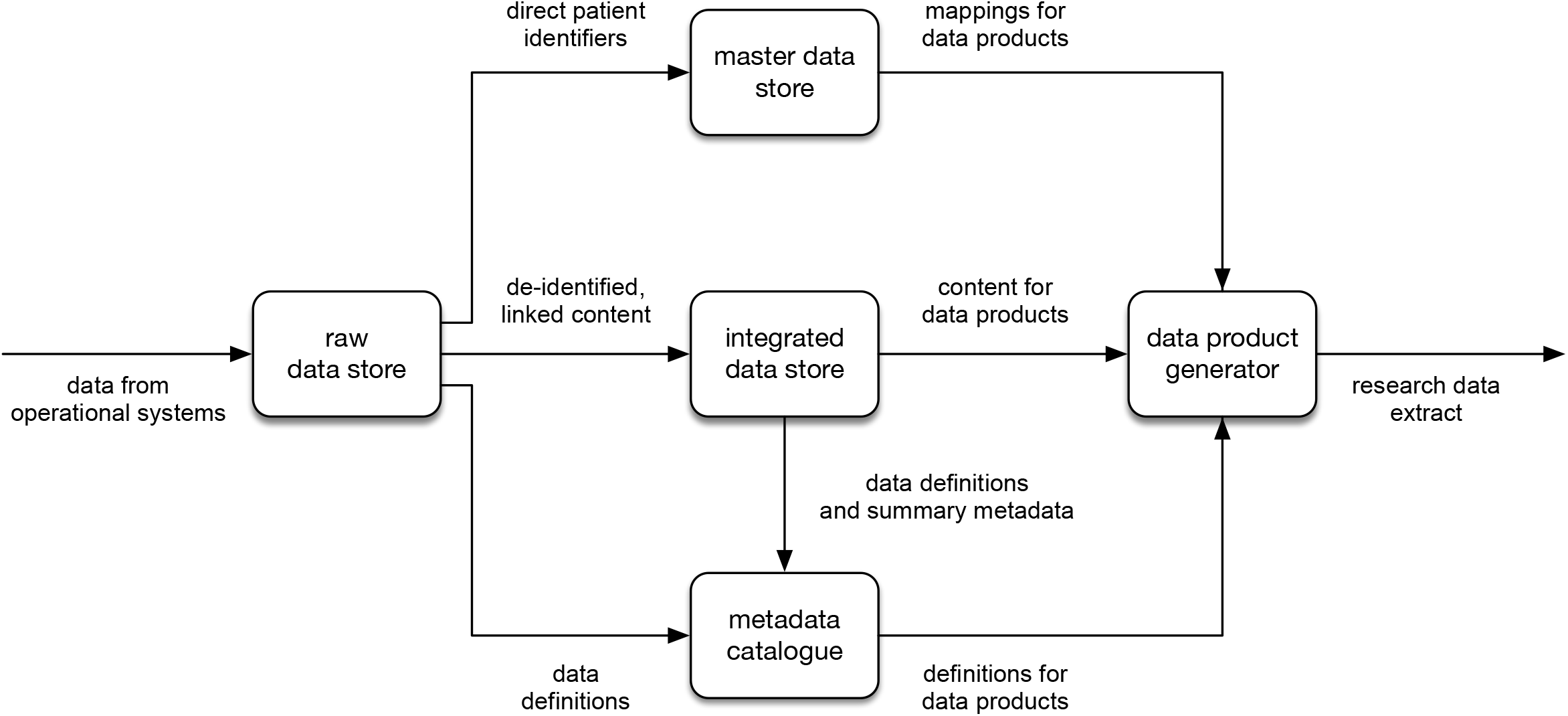
Flow diagram to depict collection, storage and output of electronic clinical data from a Health Informatics Collaborative data warehouse. The data warehouse receives data from operational systems within the hospital such as electronic patient records and laboratory information management systems (LIMS) and maps this data to individuals where the identifiers are then stored in the master data store and provides the mappings for data products. De-identified linked data is stored separately and forms the content of data products. Definitions of data items are recorded in the metadata catalogue. Data items for the data product are selected using the definitions in the metadata catalogue the mappings for these are retrieved from the master data store and data retrieved from the integrated data store to create the final data product.

**Table 5:**
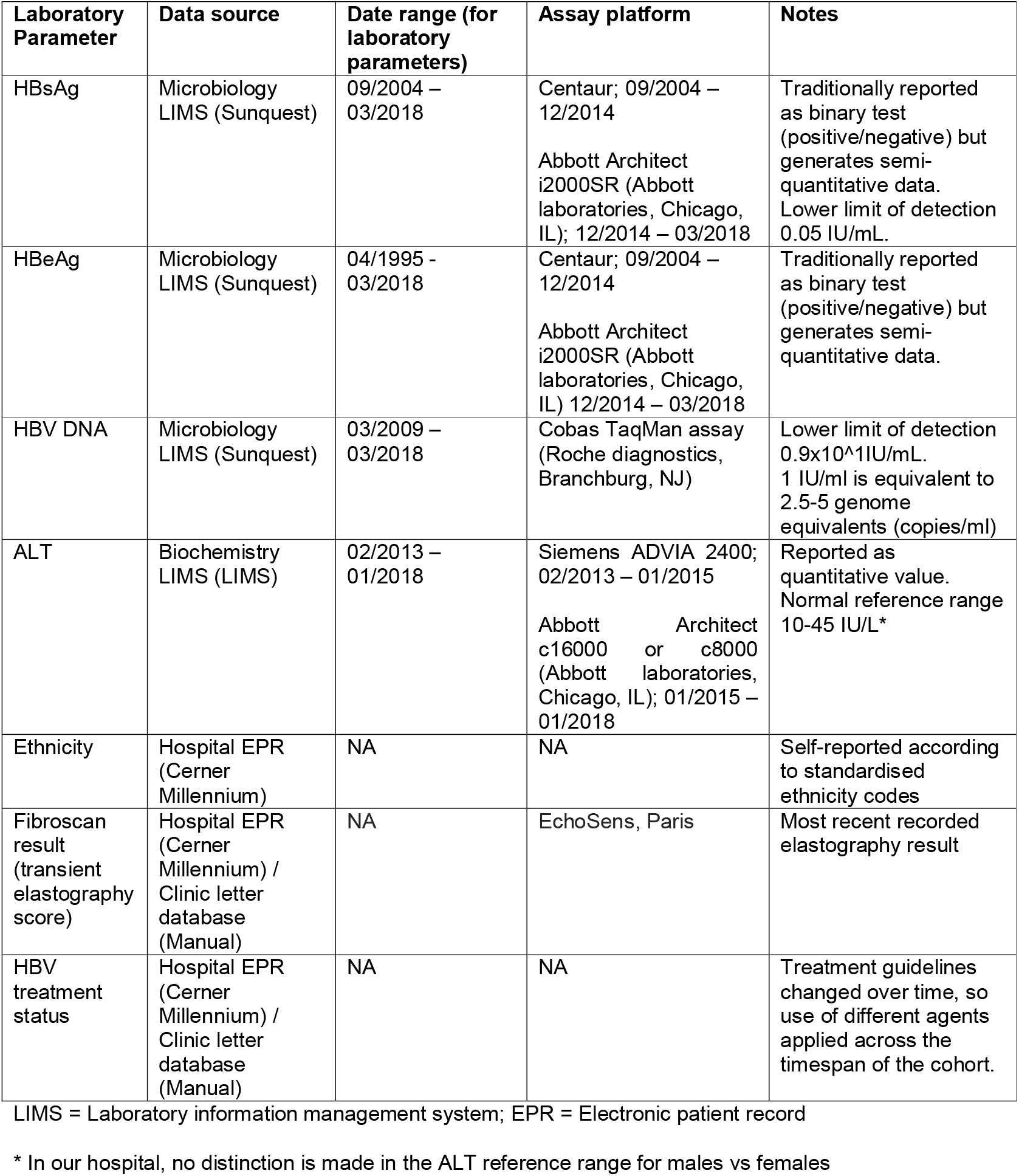
‘Data dictionary’ of clinical and demographic parameters collected for cohort of individuals with chronic HBV infection

We devised classification criteria for HBsAg and HBeAg to sort each individual into a category based on the dynamics of these serologic markers (Table 1). For HBsAg and HBeAg ‘clearers’ and HBsAg ‘potential clearers’, data which were not captured electronically or were not available from the data warehouse e.g. (most recent transient elastography score and HBV treatment status), were retrieved from the patient’s written clinical record or from dictated letters from the viral hepatitis clinic.

### Ethics

The NIHR HIC Viral Hepatitis database was approved by the NRES Committee South Central-Oxford C on 6^th^ October 2015 (REC reference: 15/SC/0523).

### Statistical analysis

We cleaned and analysed data using R and the data.table package (41). The clearance rate was 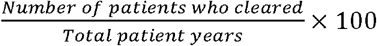. Plots were created using ggplot2 (42), and survival analysis and Kaplan-Meier plots created using the survival and rms packages (43). We used Wilcoxon or Kruskal Wallis tests for mean comparison of continuous variables, Fisher’s exact test for comparison of categorical variables, and logistic regression for multivariable analysis. We included all the parameters in our dataset in multivariable analysis, based on existing biological reasons to believe them likely to be relevant. Specifically, age at first HBsAg test is known to be associated with HBsAg clearance, sex and ethnicity could indicate diffrences in host genetics and immune response, and HBeAg status is a known marker of disease severity (20, 26, 37). Code used for this analysis is available in the attached HBsAg_Final_Analysis.html file included in the supplementary information. To define HBsAg clearance time-frames, we measured from the time of the last HBsAg result of >1000 IU/ml (or the result closest to 1000 IU/ml) to the time point at which HBsAg first became undetectable. For analysis of ALT, we used the result corresponding to the time of the first HBsAg test result.

## SUPPLEMENTARY DATA

On acceptance for publication, Supplementary Data will be made available at DOI: 10.6084/m9.figshare.7262957. Prior to publication, these files can be accessed using the following URL: https://figshare.com/s/82db3b5cd1dc5c6dd566

***Supplementary Table 1:* Summary of studies reporting rate and quantitation of HBsAg clearance in individuals with HBV infection** – Listed studies were identified from a PubMed search performed in April 2018 with the search terms: (‘hepatitis B’ OR ‘HBV’) AND (‘clearance’ OR ‘seroclearance’ OR ‘vir* negative’), written in English between 2008 – 2018, and reporting a cumulative or annual clearance rate of HBsAg using a quantitative assay.

***Supplementary Table 2:* Data for 319 adults with chronic HBV infection**

***Supplementary Figure 1:* Longitudinal data for 13 adults with chronic HBV infection who completely cleared HBsAg** – Each individual is labelled with a unique anonymised ID number, prefixed HBS. Time is shown in weeks since the first HBsAg positive test. Units (y-axis) are shown in IU/ml, except for ALT which is shown in IU/L.

***Supplementary Figure 2:* Longitudinal data for 27 adults with chronic HBV infection on a potential HBsAg clearance trajectory.** Each individual is labelled with a unique anonymised ID number, prefixed HBS. Time is shown in weeks since the first HBsAg positive test. Units (y-axis) are shown in IU/ml, except for ALT which is shown in IU/L.

***Supplementary Figure 3:*** Longitudinal data for 279 adults with chronic HBV infection who did not clear HBsAg. Each individual is labelled with a unique anonymised ID number, prefixed HBS. Time is shown in weeks since the first HBsAg positive test. Units (y-axis) are shown in IU/ml, except for ALT which is shown in IU/L.

***Supplementary Figure 4:* Boxplot showing the distribution of age among individuals who clear or potentially clear HBsAg (n=40, median age 40) and those who do not clear HBsAg (n=279; median age 34).**

***Supplementary Code:*** html file

## AUTHORS’ CONTRIBUTIONS

- Conceived the study: KJ, PCM
- Assimilated/analysed clinical records: LD, SL, JM, MP, SC, KJ
- Established the BRC informatics working group: KC
- Developed the health informatics (HIC) pipeline: DS, HS, JD, OF, KV, KW
- Lead for viral hepatitis health informatics: EB
- Analysed the data: DS, ALM, MAA, SL, JM, PCM
- Literature review: LD, JM
- Wrote the manuscript: LD, DS, PCM
- Edited and approved the final manuscript: all authors

## ACKNOWLEDGEMENTS

This work has been facilitated by infrastructure development at Oxford Biomedical Research Centre (BRC) funded by the NIHR Health Informatics Collaborative (HIC).

An earlier version of this dataset was presented as a poster at the EASL International Liver Congress meeting, Paris 2018 (https://f1000research.com/posters/7-917).

## ABBREVIATIONS

CHB: Chronic Hepatitis B virus infection
EPR: Electronic patient record
HBeAg: Hepatitis B ‘e’ antigen
HBsAg: Hepatitis B surface antigen
HBV: Hepatitis B virus
HCV: Hepatitis C virus
HIC: Health Informatics Collaborative
HIV: Human immunodeficiency virus
LFT: Liver function tests
LIMS: Laboratory information management system
NA: Nucleos(t)ide analogues
NIHR: National Institute for Health Research
PEG-IFNα: Pegylated interferon alpha 2
TDF: Tenofovir Disoproxil Fumarate
RBV: Ribavirin

