## Supplementary data table 1 and supplementary figures 1-4 for "Data from an electronic health informatics pipeline to describe clearance dynamics of Hepatitis B surface antigen (HBsAg) and e-Antigen (HBeAg) in chronic HBV infection"

**TABLES:**

**Supplementary Table 1: Summary of studies reporting rate and quantitation of HBsAg clearance in individuals with HBV infection**.........................................................................................................................2

**Supplementary Table 2: Data for 319 adults with chronic HBV infection**............see separate xls and csv file

**FIGURES:**

**Supplementary Figure 1: Longitudinal data for 13 adults with chronic HBV infection who completely cleared HBsAg**..................................................................................................................................................3

**Supplementary Figure 2: Longitudinal data for 27 adults with chronic HBV infection on a potential HBsAg clearance trajectory**.........................................................................................................................................4

**Supplementary Figure 3: Longitudinal data for 279 adults with chronic HBV infection who did not clear HBsAg**...............................................................................................................................................................5

**Supplementary Figure 4: Boxplot showing the distribution of age among individuals who clear or potentially clear HBsAg (n=40, median age 40) and those who do not clear HBsAg (n=279; median age 34)** .........................................................................................................................................................................6

**References**.......................................................................................................................................................6

***Supplementary Table 1:* Summary of studies reporting rate and quantitation of HBsAg clearance in individuals with HBV infection -** Listed studies were identified from a PubMed search performed in April 2018 with the search terms: (‘hepatitis B’ OR ‘HBV’) AND (‘clearance’ OR ‘seroclearance’ OR ‘vir* negative’), written in English between 2008 – 2018, and reporting a cumulative or annual clearance rate of HBsAg using a quantitative assay.

| **Citation (reported in alphabetical order of first author)** | **Study type and location** | **Cohort size and characteristics** | **HBsAg clearance rate** | **Factors associated with HBsAg clearance** | **Factors associated with HBsAg persistence** |
| --- | --- | --- | --- | --- | --- |
| Hara *et al*.,  (2014) (1) | Prospective study. Japan | N=553 previously NA‐naïve patients with CHB started on ETV | Cumulative clearance rate on therapy 3.5% | • HBV DNA level at baseline <3.0 log copies IU/mL  • HBsAg level <500 IU/mL | Not reported |
| Kim *et al*.,  (2014) (2) | Prospective study. Korea | N=5409 CHB patients initially treated with 3TC or ETV | Annual clearance rate 0.33% | • Baseline alanine aminotransferase (ALT) level >5 times upper limit of normal | HBeAg positivity  High HBV DNA level |
| Kobayashi *et al*.,  (2014) (3) | Prospective study. Japan | N= 2112 Japanese patients with chronic hepatitis B | Annual clearance rate 1.75% | **Untreated:**  •Median age ≥50 years  •HBsAg ≤2,000 IU/mL  **Treated:**  •No family history  •Interferon treatment  •HBeAg-negative | Not reported |
| Kuo *et al*.,  (2015) (4) | Prospective study. Taiwan | N=59 HBsAg carriers | Annual clearance rate 2.7% | •Lower levels of HBsAg and HBV DNA | Not reported |
| Lim *et al*.,  (2016) (5) | Retrospective study. New Zealand | N=572 CHB patients, followed up for 28 years. | Annual clearance rate 1.1% | •Increasing age  •Lower baseline HBsAg levels  •Lower baseline HBV DNA levels  •Accelerated HBsAg loss in those with lower HBsAg levels | Not reported |
| Nagaoka *et al*.,  (2015) (6) | Retrospective study. Japan | N=392 Japanese CHB patients | Annual clearance rate 0.91% | •HBsAg <3.3 log IU/ml  •Treatment with NA  •Hepatic flares promoted rapid declines and greater annual reductions of HBsAg levels in patients with HBsAg clearance. | Not reported |
| Park *et al*.,  (2016) (7) | Retrospective study. South Korea | N= 1919 HBsAg carriers | Annual clearance rate 0.76% | •Not reported | Treatment exposure  Baseline HBV DNA levels of >2000 IU/ml |
| Ungtrakul *et al*.,  (2017) (8) | Retrospective study. Thailand | N=300 HBeAg-negative CHB patients with initial serum HBV/DNA levels <2000 IU/ml. | Cumulative clearance 5% after 24 months | •Quantitative HBsAg level at a single timepoint (in HBeAg-negative CHB) with low HBV DNA levels at baseline predicted spontaneous HBsAg clearance at 24 months. | Not reported |
| Jeng *et al*.,  (2017) (9) | Prospective study. Taiwan | N=1075 HBeAg negative patients treated with NAs | Annual clearance rate: on therapy 0.15%;  off therapy 1.78%. | •Shorter time to undetectable HBV DNA (<12 weeks)  •Greater HBsAg reduction during therapy (> 1 log 10) | Not reported |

CHB = chronic hepatitis B virus; NA = nucleos(t)ide analogue; 3TC = lamivudine; ETV = entecavir

***Supplementary Figure 1:* Longitudinal data for 13 adults with chronic HBV infection who completely cleared HBsAg -** Each individual is labelled with a unique anonymised ID number, prefixed HBS. Time is shown in weeks since the first HBsAg positive test. Units (y-axis) are shown in IU/ml, except for ALT which is shown in IU/L.

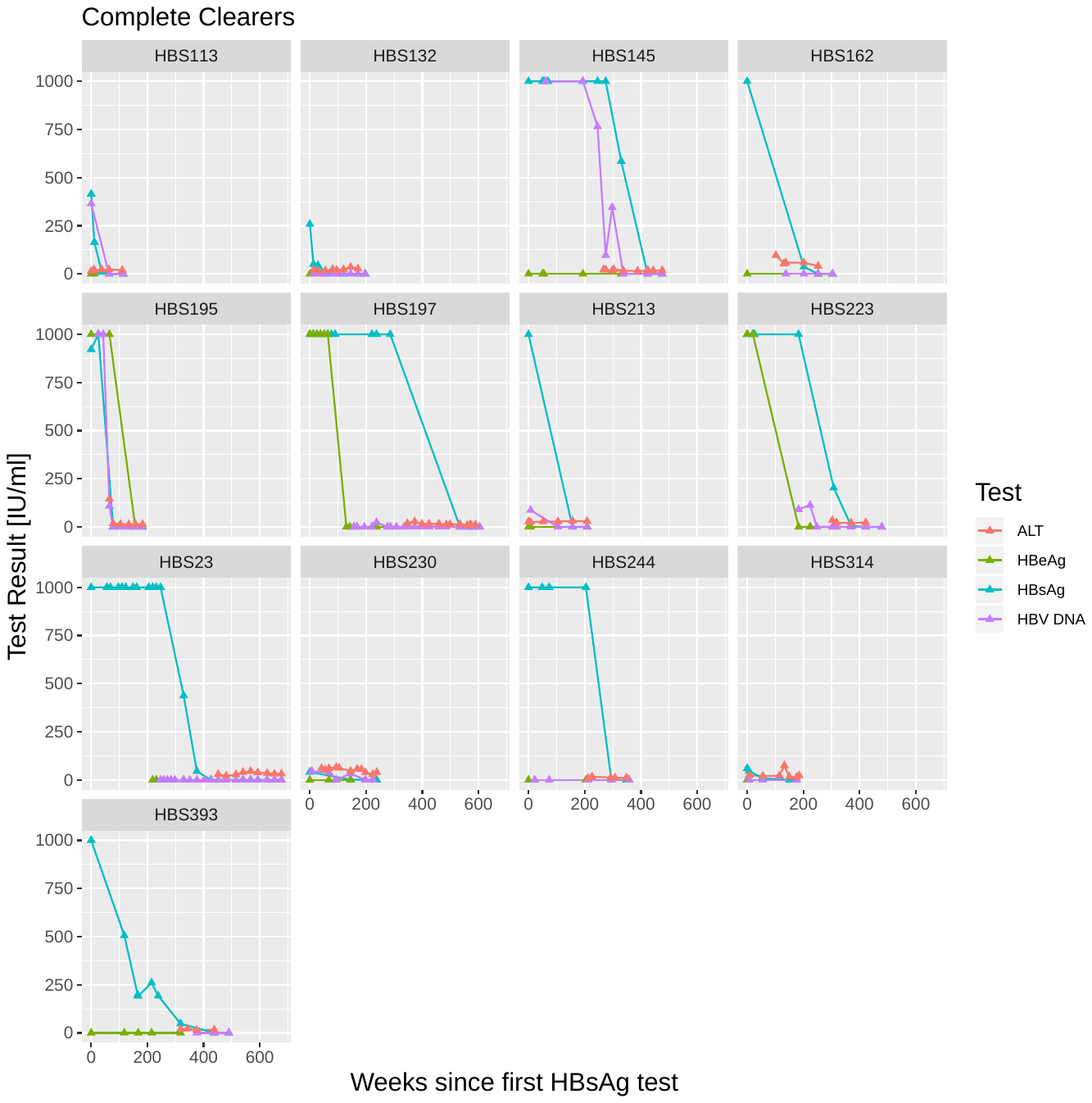

***Supplementary Figure 2:* Longitudinal data for 27 adults with chronic HBV infection on a potential HBsAg clearance trajectory.** Each individual is labelled with a unique anonymised ID number, prefixed HBS. Time is shown in weeks since the first HBsAg positive test. Units (y-axis) are shown in IU/ml, except for ALT which is shown in IU/L.

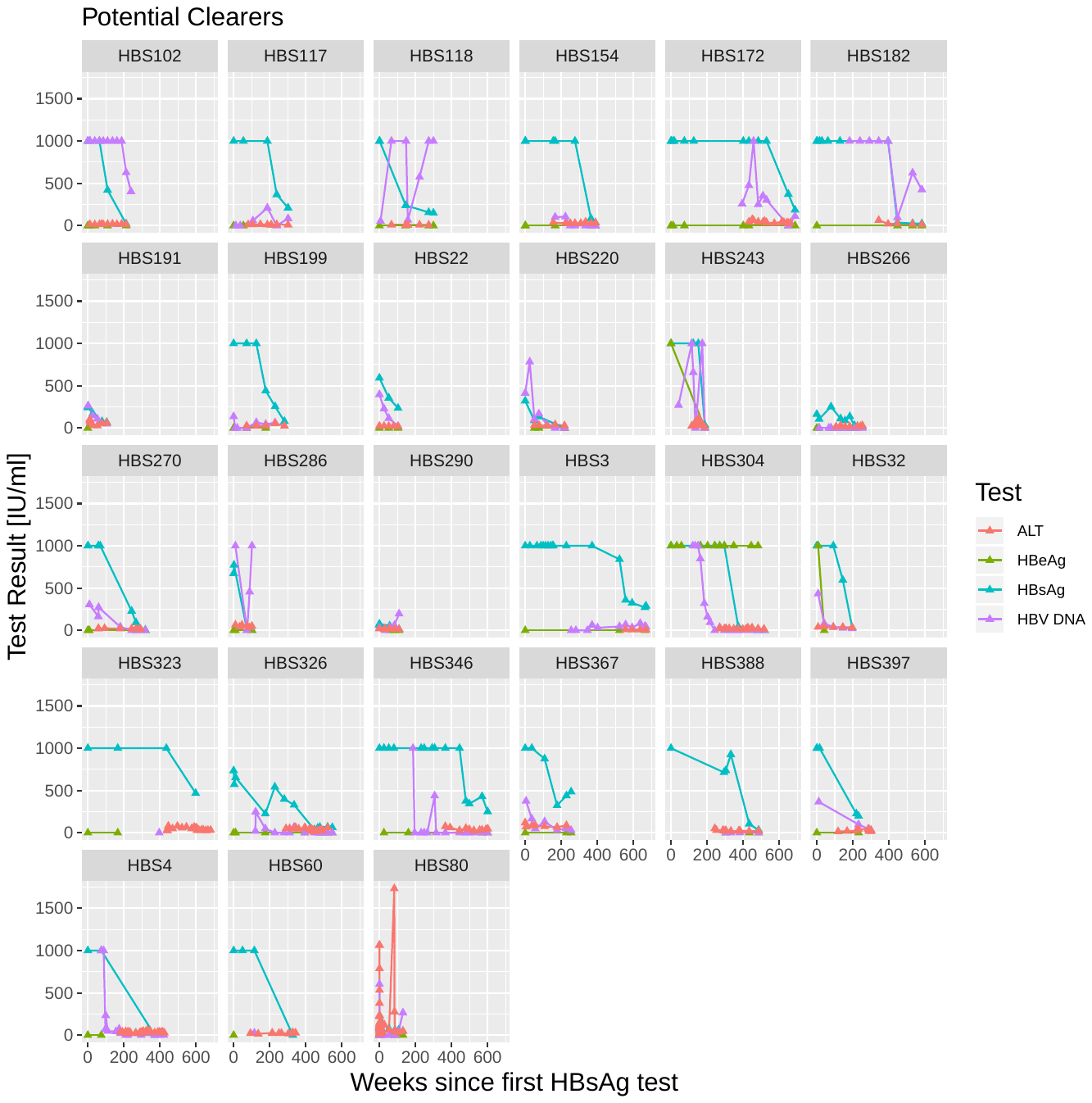

***Supplementary Figure 3:* Longitudinal data for 279 adults with chronic HBV infection who did not clear HBsAg**. Each individual is labelled with a unique anonymised ID number, prefixed HBS. Time is shown in weeks since the first HBsAg positive test. Units (y-axis) are shown in IU/ml, except for ALT which is shown in IU/L.

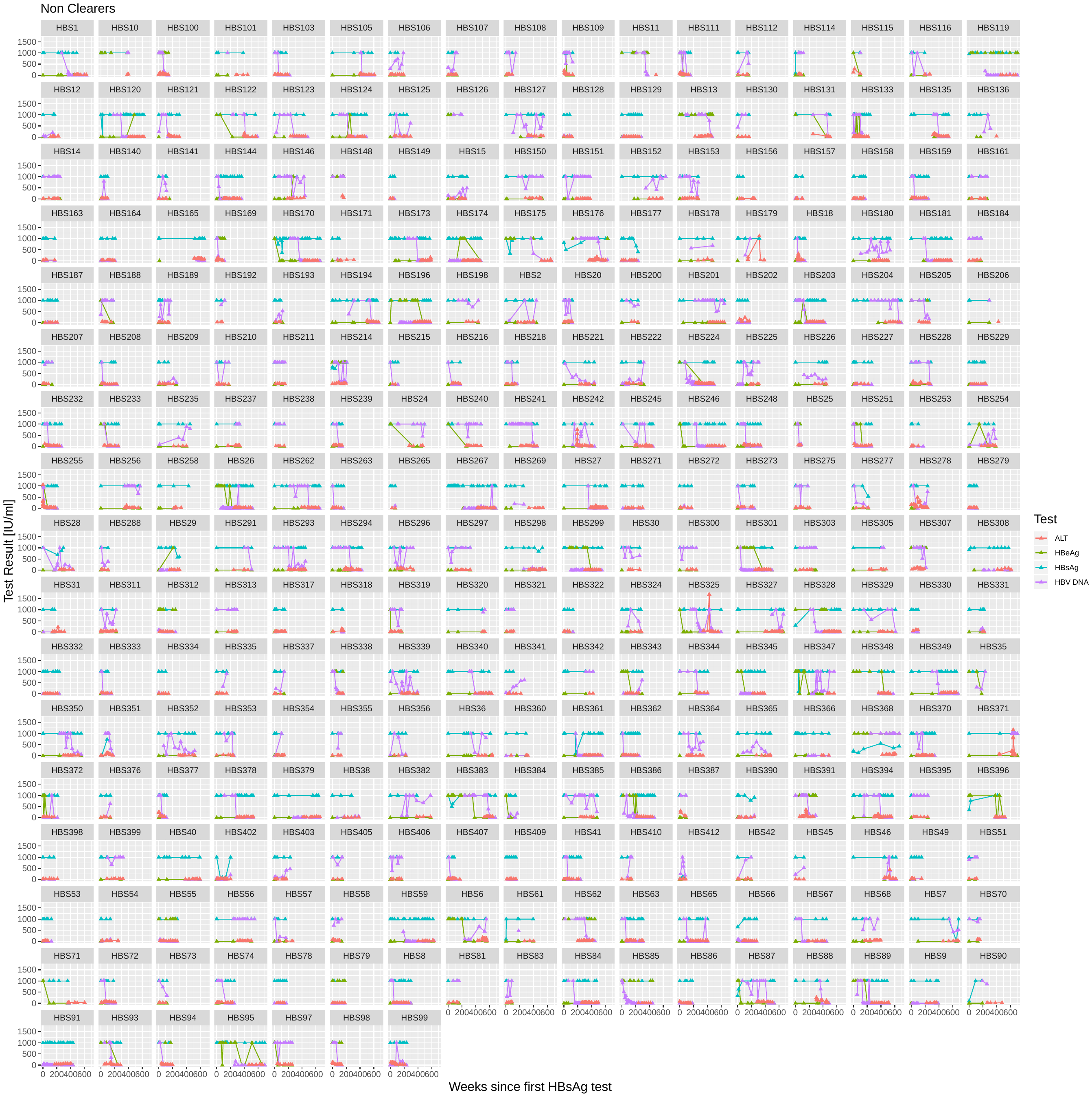

***Supplementary Figure 4:* Boxplot showing the distribution of age among individuals who clear or potentially clear HBsAg (n=40, median age 40) and those who do not clear HBsAg (n=279; median age 34).**

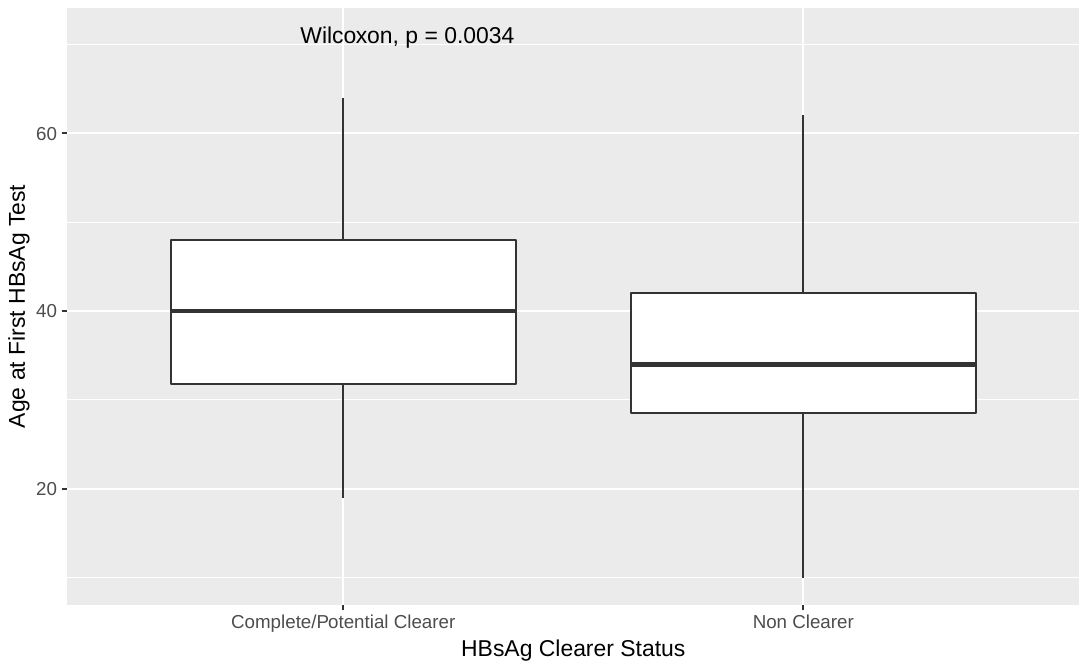
